# Sex-specific sub-lethal effects of low virulence entomopathogenic fungi may boost the Sterile Insect Technique

**DOI:** 10.1101/2024.03.07.583916

**Authors:** David Duneau, Romain Gallet, Maureen Adhiambo, Emilie Delétré, Anais Chailleux, Fathiya Khamis, Sevgan Subramanian, Thierry Brévault, Simon Fellous

## Abstract

**BACKGROUND:** The Sterile Insect Technique (SIT) is a species-specific method for controlling insect pests. Recent studies have explored the combination of SIT with entomopathogenic microorganisms, known as boosted-SIT, to enhance its effectiveness. This study aimed to evaluate the potential of the entomopathogenic fungi, *Metarhizium anisopliae*, in boosting the SIT for managing the oriental fruit fly, *Bactrocera dorsalis*.

**METHODS:** Adult flies from a laboratory population of *B. dorsalis* were inoculated with one of eight strains of *M. anisopliae* to assess fungus virulence in each sex. Ideally, boosted-SIT should minimally impact sterile males and reduce female fecundity maximally. A brief exposure to fungal spores was efficient to infect males, and for them to transmit the fungus to females when hosted together for 24 hours.

**RESULTS:** Our results showed significant variations in the mortality induced by the strains in males, but not in females that exhibited low mortality. Strains varied in their sub-lethal effects on female fecundity, with almost a two-fold variation among strains. Furthermore, strains that had the lowest virulence on males tended to reduce female fecundity the most.

**CONCLUSION:** Our study brings a proof of concept that it is possible to leverage boosted- SIT using carefully selected pathogen strains and their sub-lethal effects on both the male and female fruit fly.

## Introduction

Society’s expectation for new crop-protection solutions is to minimize their unintended impacts of existing pest management practices on the environment and human health (see the United Nations Sustainable Development Goals to be achieved by 2030, especially SDG2 and SDG12). The Sterile Insect Technique (SIT) offers a species-specific approach to controlling insect pests. It involves using irradiation, such as gamma rays or X-rays, to sterilize mass- reared insects. These sterilized insects, while still sexually competitive, are unable to produce offspring. When released in large numbers, they reduce mating with fertile wild counterparts. If wild females mate with significantly high proportion of sterile males, the target insect population can decline and possibly collapse. Successful control using the SIT has been reported for various insects including the New World screwworm, tsetse fly, melon fruit fly, Queensland fruit fly, and pink bollworm (Dyck et al., 2021). Despite these successes, continuous research has been carried-out since SIT inception to gain efficacy and reduce costs.

Boosted-SIT, which focuses on supplementing mortality factors in addition to inducing sterility, is a promising evolution that still necessitates investigation before actual deployment. It utilizes released SIT insects to deliver biocontrol agents or chemicals to target insects that they contact. For example, in the case of tsetse flies, sterilized males can be impregnated with pyriproxyfen, an insect growth regulator molecule that inhibits larval development (Sullivan and Goh, 2008). Laboratory studies have shown that these males can transfer this insect growth regulator to females for up to ten days through simple contact, even if mating fails, effectively preventing them from producing viable offspring (Laroche et al., 2020). Mathematical and agent-based modelling identified the conditions of where field efficacy may be greater than for classical SIT (Diouf et al., 2022; Haramboure et al., 2020; Pleydell and Bouyer, 2019). Models suggest large dependence on the biological features of the host species and the type of mortality agent employed. Field experiments have been conducted testing boosted-SIT on the Mediterranean fruit fly, *Ceratitis capitata* in Guatemala. In these experiments, field-released males were inoculated with the entomopathogenic fungi, *Beauveria bassiana*, leading to the successful transmission of fungal spores to wild *C. capitata* females (Flores et al., 2013). However, inoculating pathogens in sterile males can be costly and may increase their mortality or reduce their competitiveness with wild males in the field (Flores et al., 2013; Thaochan and Ngampongsai, 2015). Therefore, considering the effects of biocontrol agents and their dosages on the survival and biological traits of male target insects is crucial for the success of boosted-SIT.

To be effective in a boosted-SIT strategy, a pathogen must possess combinations of phenotypic features that maximize deleterious effects on females of wild populations and minimize effects on released sterile male hosts. The released insects are most often males, whereas wild females are usually targeted. A promising avenue may therefore will be to select appropriate types and strains of pathogen that are associated with sex-specific infectivity variation on released insects. The insect pathology literature is rich with examples of genetic variation among strains of entomopathogenic fungi (Barzanti et al., 2023; Gasmi et al., 2021; Serna-Domínguez et al., 2019). Moreover, pathogens often have different effects on male and female hosts of the same species (Duneau and Ebert, 2012; Duneau et al., 2012, 2017; Fellous and Koella, 2009; Gipson and Hall, 2018; Zuk and McKean, 1996). Hence, our goal was to identifying sex-specific optimal pathogenic strains and explore its potential to enhance the effectiveness of boosted-SIT programs.

We tested the idea of exploiting inter-strain variability to boost boosted-SIT with the invasive Oriental fruit fly, *Bactrocera dorsalis*. It is recognized as one of the most destructive pests in fruit production that is rapidly spreading across the world (Drew et al., 2005; Mutamiswa et al., 2021; Nugnes et al., 2018). It has become a major threat to mango orchards and has prompted numerous control efforts (Ekesi et al., 2016a, 2016b). Among the available control methods, SIT has proven to be effective (Orankanok et al., 2008), and the independent use of the entomopathogenic fungus *Metarhizium* is widely deployed (Faye et al., 2021; Gichuhi et al., 2020; Sookar et al., 2014). Spores of *M. anisopliae* can be horizontally transferred from males to females during sexual contact as demonstrated in the closely related species *Ceratitis capitata* (Dimbi et al., 2013). Isolates of *Metarhizium* species also exhibit varying degrees of pathogenicity towards *Bactrocera tryoni* (Carswell et al., 1998; McGuire et al., 2023). Screening studies have revealed that different isolates of *M. anisopliae* can induce varying levels of mortality in various insect species (Akutse et al., 2020; Peng et al., 2022), including tephritid flies (Moraga et al., 2006), and specifically *B. dorsalis* (Aemprapa, 2007). However, while most studies have focused on identifying the most virulent strains with the highest mortality rates, our approach took a different perspective.

We aimed to identify a strain that minimally affects male flies while imposing a significant cost on female flies, thereby enhancing the overall success of the control method. To achieve this, we tested eight *M. anisopliae* strains on the males and females of a laboratory population of *B. dorsalis*. Such an approach can maintain the competitiveness of male fruit flies to access to the females, while contributing to horizontal transmission of *M. anisopliae* which increases female mortality. Our results indicated significant variations among the strains, suggesting that certain strains may be more effective than others in achieving successful control measures considering sex-specific sub-lethal effects. Based on our findings, we can make a recommendation regarding the selection of fungal strains for implementing boosted-SIT.

## Material and Methods

### Biological material

We used the laboratory population of *B. dorsalis* maintained and produced by the Animal Rearing and Containment Unit (ARCU) of the International Centre for Insect Physiology and Ecology (*icipe*) in Nairobi, Kenya. This population originated from insects that emerged from infested mango fruits purchased at a local market in Nairobi, Kenya. Fly production was performed according to the methods outlined by Ekesi et al. (2007). They were kept in stable conditions of 28 ± 1°C with a relative humidity (RH) of 50 ± 8% and a photoperiod of L12:D12. When experiments began, the colony had over a thousand individuals and was over 100 generations old. However, the introduction of wild flies once or twice a year brought new alleles in a population otherwise selected to live in laboratory conditions and most likely with a smaller genetic diversity. The strains of *M. anisopliae* used for this experiment are described in table 1; all belonged to *icipe*’s Arthropod Pathology Unit Germplasm (Entomopathogens collection). Fungal conidia (i.e., spores hereafter), the infective stage, were produced by growing the fungi on sterilized rice in Petri dishes. These strains were chosen based on their previously demonstrated infectivity and lethality against *B. dorsalis* (Ouna et al., 2010).

**Table 1:** *M. anisopliae* strains used in the study.

| Strain name | Substrate or host & Country | Year of isolation |
| --- | --- | --- |
| ICIPE 402 | Homoptera, Kenya | 2007 |
| ICIPE 7 | *Amblyomma variegatum*, Kenya | 1996 |
| ICIPE 41 | Soil, Kenya | 1990 |
| ICIPE 18 | Soil, Kenya | 1989 |
| ICIPE 95 | Sand fly, Kenya | 2005 |
| ICIPE 387 | *Forficula senegalensis*, Kenya | 2007 |
| ICIPE 55 | Soil, Kenya | 2005 |
| ICIPE 69 | Soil, Democratic Republic of | 1990 |
|  | the Congo |  |

### Methods

Groups of 13 males were inoculated in custom-made inoculators, a 98 mm by 45 mm plastic cylinder with velvet on the side and nylon netting on the two ends (Dimbi et al., 2013). Each inoculator received 0.3 g of dry spores that were evenly spread on the velvet. Male flies (n=10) remained in the inoculator for three minutes to facilitate pick up of spores before either being used to estimate inoculum size (n = 3) or being dispatched in a cage that contained clean non inoculated female flies (n=20). The number of spores was estimated using a Malassez cell counting chamber after grinding each individual male in 1 mL of water. Each experiment was replicated three times for each fungal strain.

Each assay included exposing 20 virgin females to 10 inoculated-males in a 20 cm cubic netting-cage. The cages were equipped with a water-soaked cotton ball and a dish containing *ad libitum* yeast extract and glucose powders, each provided separately. Both the cotton ball and the dish were replaced daily to provide the flies with unrestricted access to food and water. Additionally, the cages included the skin of one (replicated experiment 1 and 2) or two (replicated experiment 3) half mango (i.e., dome), devoid of its flesh, which served as an oviposition substrate.

Males and females were housed together in the same cage for a duration of 24 hours, allowing them to mate. Following this initial period where no death occurred, males and females were separated into individual cages. Daily, any flies that died were tallied and promptly removed. Mango skins were changed every day and egg number were counted. The experiments were replicated three times per block on different days. Each block contained unexposed males (control) as well as males exposed to the fungal strains.

### Statistical analyses

All analyses were performed using *R v4.1.0* and *Rstudio* (R Core Team, 2020; RStudio Team, 2016). Supplementary material 1 was generated by *Rmarkdown* (Allaire et al., 2023), a component of RStudio, and provided a summary data table, with all scripts and their associated analyses and plots, including supplementary figures. All the analysis and illustrations were done with the *tidyverse v1.3.2* R package suite (Wickham et al. 2019).

#### Spore inoculum

To test for differences in spore inoculum between fungal strains we fitted a linear model with the function *fitme* from the package *spaMM v4.0.0* (Rousset and Ferdy, 2014). We then estimated the 95 % confidence intervals of the log2 of the spore inoculum for each strain based on boostraps done with the function *confint* from the same package. We confirmed the parametric graphical approach of the differences using a Kruskal-Wallis test, comparing all the strains in the same test.

#### Survival

To examine the differences in baseline mortality between male and female flies and across different experiment days, we used a semi-parametric Cox model using the *coxph* from the package *survival v3.2.11* (Therneau and Grambsch, 2000). Using the R synthax the model was specified as follow: *coxph(Surv(Time_post_inoculum, Censor)∼ Sex * Day of experiment + (1|Cage_ID)*. We considered that flies were in different cages by including the factor Cage_ID as a random effect. To assess the disparities in survival among strains for males, following spore inoculation, and for females, following contact with males, we employed a single Cox model encompassing both sexes and all strains: *coxph(Surv(Time_post_inoculum, Censor)∼ Fungal_treatment + Fungal_treatment : Sex + (1|Cage_ID)*. This allowed us to extract the log hazard ratio for each strain relative to the most virulent strain (ICIPE 69). We considered that flies were in different cages as in the previous model. Since strains were tested on different days without replication over multiple days and only two to three strains were evaluated per day, we did not include the “day of experiment” as a factor in the model. This was supported by the fact that baseline mortality was low and did not differ across days. The *control* treatment was analyzed separately and not included in this particular model, as we focused on comparing strains, which is a more conservative approach. All these statistical decisions were carefully considered in our interpretation of the results. Significant differences were determined by the overlap of the 95 % confidence interval, detailed p value in comparison to ICIPE 69 can be found in supplementary material.

#### Fecundity

To assess the differences in egg numbers among cages of females that had been in contact with males inoculated with different fungal strains, we used the Kruskal-Wallis test, followed by Dunn posthoc tests conducted separately for each experiment day. This approach was chosen due to the non-linear nature of daily egg numbers, the presence of three replicate cages per strain, and the high repeatability of egg counts among cages. By using these methods, we ensured a conservative approach to evaluate the variations in egg numbers between different treatments.

## Results

*Spore inoculum.* We first tested whether spores were properly inoculated to male hosts with our approach and whether all strains were similarly inoculated. Males were inoculated with approximately 250-1000 spores (Figure 1). The 95% confidence intervals of the average inoculum for all strains overlapped with each other, supporting that there were no significant differences among the strains for the inoculum (Kruskal-Wallis test: df = 7, χ^2^ = 11.4, p = 0.12).

**Figure 1:**
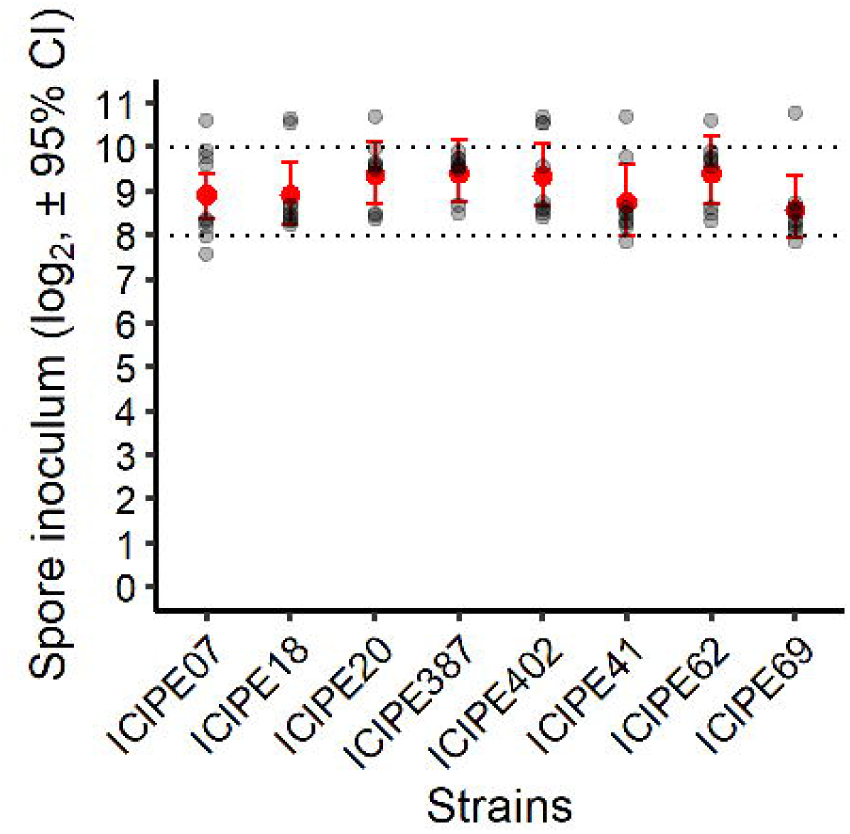
Spore inoculum for each fungal strains. Males were inoculated with approximately 250-1000 spores (i.e., between 2^8 and 2^10 spores). The 95 % confidence intervals for all strains overlapped with each other, suggesting that there are no significant differences among the strains for the inoculum. Red dots and error bars represent the means of spore inoculum and their 95 % confidence intervals estimated by bootstraps.

### Fly survival to infection with various fungal strains

We monitored the survival of male and female flies for a duration of 7 days. There was no significant difference in baseline mortality, observed when individuals were not exposed to spores, between sexes and across different exposure days (P-value from cox model: Sex = 0.07; Day of inoculum = 0.3, and Interaction = 0.96, details in supplementary material 1, section 2).

Males were exposed to spores from various strains (n = 7) through inoculation, while females were exposed through contact with these males. Both male and female individuals exhibited clear mortality following the infection (Figure 2A). Females experienced mortality approximately 1 day after males, suggesting a lag associated with transmission from males to females. Males, who were directly inoculated with spores, experienced higher mortality rates from the infection compared to females exposed to infected males through contact. All male individuals succumbed to the infection 6 days after inoculation, whereas more than 50 % of the females remain alive even after 7 days from the start of the experiments. Nevertheless, we cannot exclude at this point that females which survived were unexposed to the spores.

**Figure 2:**
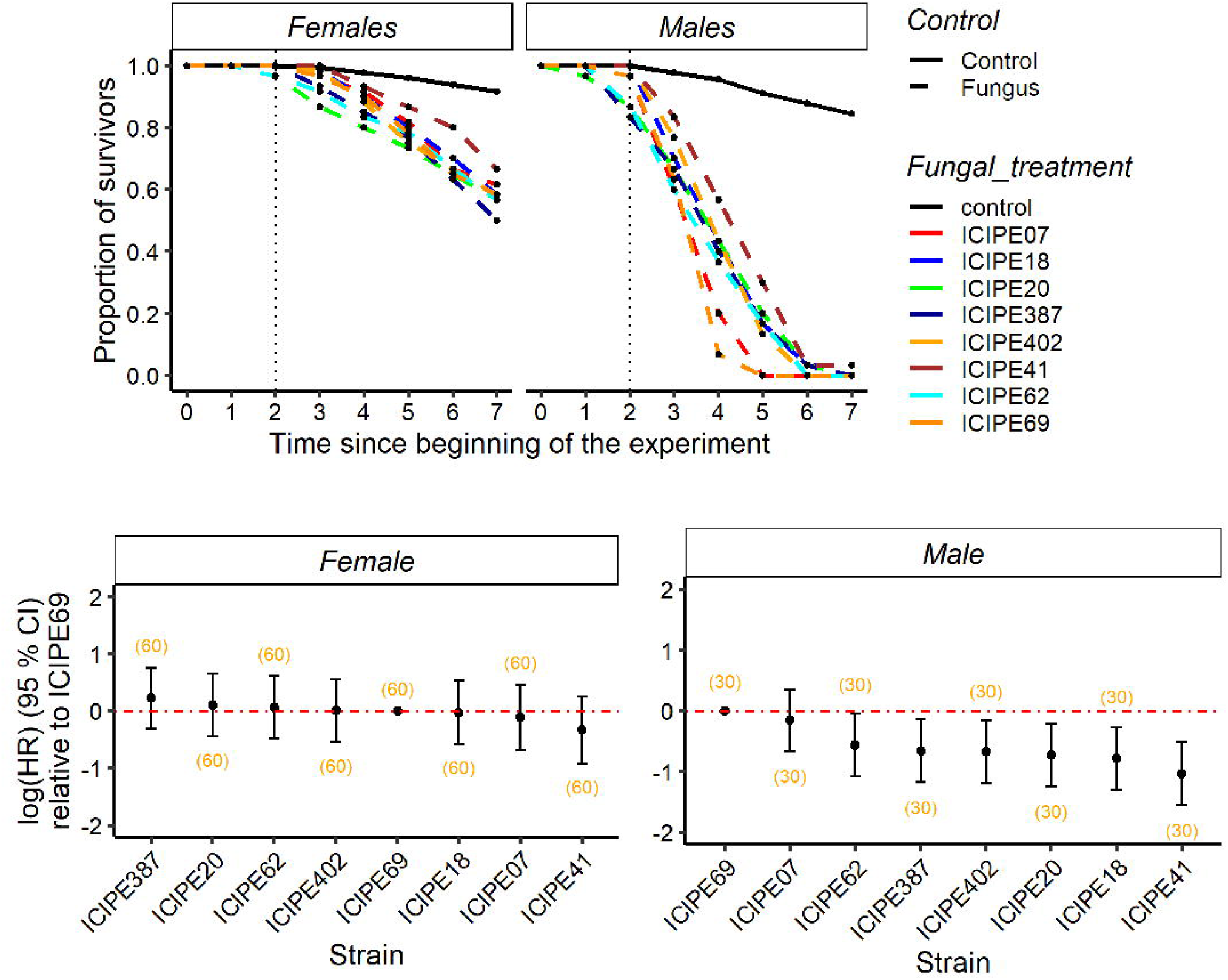
Survival of male and female oriental fruit flies across eight strains of the pathogenic fungus Metarhizium anisopliae. A- Proportion of survivors over time. Males were inoculated, and females were exposed to the inoculated males for 24 hours. After separating males and females into cages, survival rates were recorded daily. B- Relative difference in risk of mortality. The risk of dying, represented by the log(Hazard Ratio), was calculated for each strain based on the survival rates mentioned in A. Positive values indicate a higher risk of mortality. Error bars represent the 95 % confidence interval. The red line represents the reference strain (ICIPE69). Statistical significance is indicated when the error bars do not overlap or cross the red line. Sample sizes from three replicated experiments are indicated in brackets.

There was no significant difference in female mortality between strains (Figure 2B). However, strains ICIPE 7 and ICIPE 69 exhibited a higher rate of male mortality compared to other strains, especially compared to the least pathogenic strain ICIPE 41 (Figure 2B). It is important to note, however, that the most virulent strains were tested on the same day (Supplementary figure 1). While the baseline mortality on that day was not higher, it is worth considering that the higher mortality could potentially be influenced by environmental conditions specific to that day.

As anticipated due to the lack of statistical differences in spore inoculum among strains, there was no correlation observed between the inoculum to males and their survival (Pearson correlation test: df = 6, cor = -0.33, p = 0.4), suggesting that the difference in virulence is due to the pathogenicity of the strains not to the inoculum (supplementary figure and statistical details in Supplementary material 1, section 2).

### Impact of fungal transmission on female fecundity

During the 7-day duration of the experiment, we assessed the fecundity of female flies on mango skin. We observed a significant difference in fecundity among treatments (Figure 3A, statistical details in Supplementary material 1). In general, most strains resulted in a decrease of the number of eggs per cage, indicating a potential impact on fecundity. Interestingly, strains ICIPE 69, and ICIPE 7 exhibited contrasting behaviour, as they showed higher fecundity compared to the control despite their high virulence (Figure 3A). What were the drivers of these fecundity variations? At the beginning of the experiment, each cage contained 20 females, and their numbers gradually decreased throughout the study period as they die. As a result, the egg count was not based on a constant number of females, and the reduction of fecundity per cage could be due to female mortality within the cages. We tested the difference of egg numbers per cage within the 2 days after females were in contact with males as there was no mortality at this stage. The pattern did not differ strongly with the total number of eggs per cage over a span of 7 days, suggesting that the difference in number of eggs was not only driven by the reduction in number of females per cage over time but also by the impact of the fungus directly on female fecundity (supplementary figure and statistical details in Supplementary material 1, section 3).

**Figure 3:**
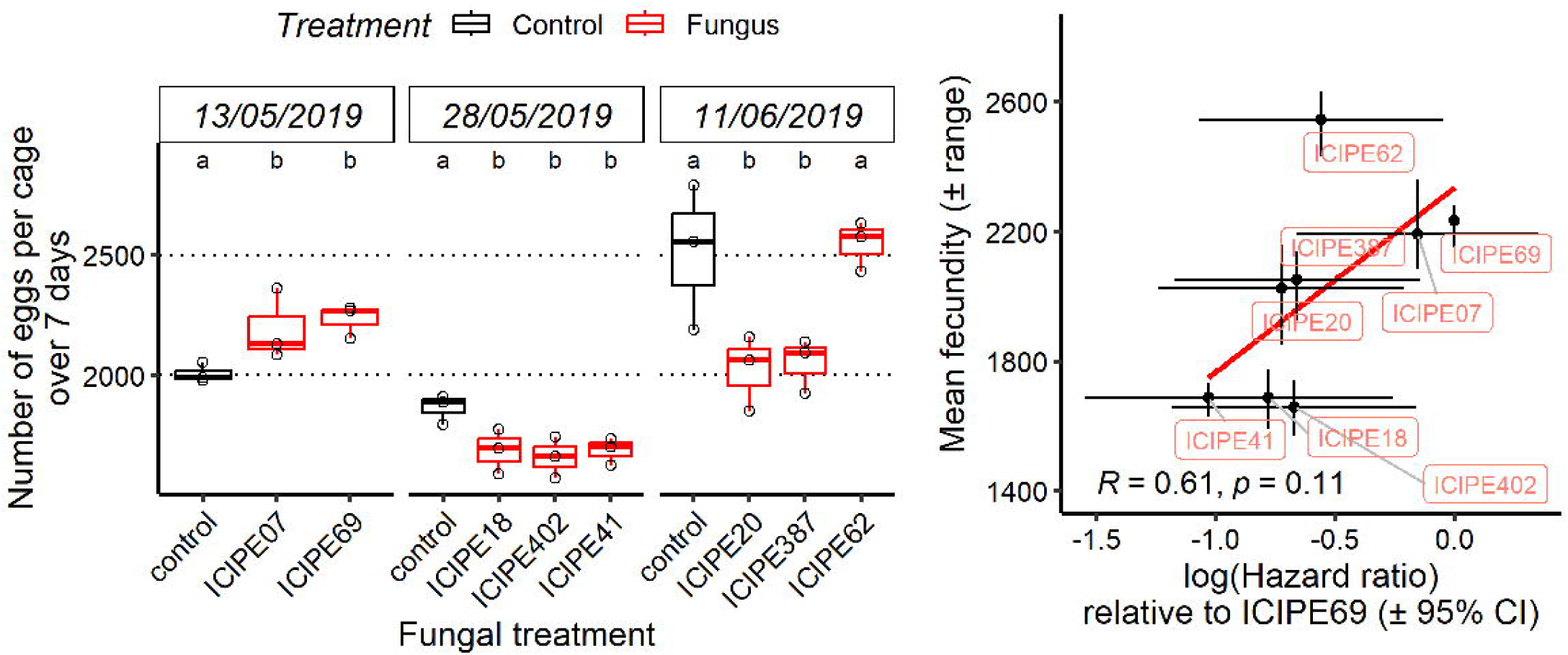
Impact of fungal treatment on female fecundity. A- Total number of eggs per cage. The total number of eggs laid within a 7-day period was recorded for each of the eight fungal strains. Boxplots represent the 25th, 50th (median), and 75th percentiles. Due to our experimental design with a limited sample size of three cages per treatment and the use of non-parametric statistical tests, the letters in the figure represent the statistical differences after p-value corrections with the probability to falsely attribute a difference (error type I) of 10%, instead of 5 %. B- Correlation between severity in males (lethality) and severity in females (fecundity). There is evidence supporting the observation that strains causing slower mortality in males also result in a greater reduction in female fecundity. The correlation and the pvalue were the results of a Pearson correlation test.

We analysed the correlation between survival (log(HR)) and fecundity (mean) and, despite a limited number of strains, we observed a tendency where strains with lower mortality rates also exhibited greater reductions in fecundity (Figure3B, Pearson correlation = 0.6, p-value = 0.1).

## Discussion

Our experiments investigated the inter-strain variability of a fungal pathogen to identify trait combinations in each host sex able to improve the efficacy of the boosted-SIT. To this end, we mimicked the release of sterile males inoculated with *M. anisoplae* spores in populations of *B. dorsalis* flies. We observed that among the eight tested strains, there was variability in their virulence on males. Females did not die strongly after interacting with these males, and the virulence in females did not significantly differ across the different pathogen strains. However, female egg production, which directly determines damages to fruit crops and the dynamics of their populations, depended on the inoculated strain.

Our study successfully demonstrated that the inter-strain diversity of a fungal pathogen could be relevant when designing a boosted-SIT program using live pathogens. In particular, we observed substantial variations among strains in terms of host fecundity following fungal-exposure. The control of insect reproduction, and the limitation of its access to valuable crops, are hallmarks of contemporary pest management programs (Dyck et al., 2021). This approach contrasts with insecticide-based strategies that usually rely on killing as many insects as possible. In the present case, reducing wild female fecundity may be most beneficial when this parameter determines the fate of the population. This could be particularly the case when targeted females mate multiple times, use sperm acquired before the release of sterile males, or when the ratio of sterile to fertile males is low (Dyck et al., 2021). Because each insect species has specific biological features, the most useful combinations of fungal traits are bound to depend on factors such as host and pathogen species and the ecological context. Any boosted-SIT program should hence include a preliminary analysis of the desired effects of the bioagent on rates of transmission, latency in both sexes, mortality and fecundity of females.

Two alternative methods of Boosted-SIT have been proposed: using live pathogens or chemical insecticides. In both cases, diminishing health costs for the inoculated insects, which are released and must interact with wild insects, is paramount. When using insecticides, some authors suggested using molecules with life-stage specific effects. Pyriproxifen is one such insecticide, it is only toxic to eggs and larvae and minute amounts suffice to exert juvenile mortality (Hustedt et al., 2020). When using live pathogens, a challenge emerges from so-called dose effects. Infectious dose, the number of parasite propagules hosts are exposed to, generally determines the outcome of infection, sometimes in combination with other parameters such as host physiology, sex or environment (Ben-Ami et al., 2008; Fellous and Koella, 2010; Lunn et al., 2019). In simple terms, the greater the dose, the greater the virulence. In the case of boosted-SIT, released sterile males necessarily receive larger doses than wild female insects, as only a fraction of the inoculated propagules are transmitted during field interactions. This could pose a problem when aiming to use a dose that have a minimal impact on the sterile males but maximal on the females.

Host sex is a factor known to determine both pathogen virulence and transmission (Duneau and Ebert, 2012; Duneau et al., 2012, 2017; Fellous and Koella, 2009; Gipson and Hall, 2018; Zuk and McKean, 1996). In our study, virulence in females was relatively low compared to males. However, the infection processes were different between males and females by design, mimicking the process in boosted-SIT, which prevents us to state that females were intrinsically less susceptible than males. The difference was most probably due to a lower infectious dose in females, which could have not only reduced the observed virulence but also hampered the detection of among strain variation. Hence, our study suggests that it might be difficult to kill females with a pathogen while having reduced impact on the males, unless the difference in susceptibility between sex counters balance the difference in dose.

The alternative is if the pathogen impact female fecundity, or fertility as the chemical Pyriproxifen does (Pleydell and Bouyer, 2019). Our study suggests that indeed, although our tested pathogens were sublethal, they reduced the amount of offsprings females exposed to infected males could produce. This has strong consequences especially when sterile males are not only partially sterile but also when females can mate several times, including with wild males. Furthermore, our results suggest a correlation between pathogen effect on fecundity on females and mortality in males. Potential links between traits expressed by various pathogen strains confirm the critical need for careful selection of pathogen in Boosted-SIT. The most virulent strain may not always be the best choice, not only because it kills inoculated sterile males, but also because it could have unaccounted for effects on female fecundity and fertility.

To conclude, if boosted-SIT using live pathogens does not appear as a miracle improvement over classical SIT, refining the choice of pathogen used, based on a good understanding of the sex-specific effects needed to reduce agricultural damages, may open new perspectives. Here, we contrasted pathogen effects on mortality with those on female fecundity, revealing a potential lever to reduce fecundity in species where females mate numerous times. Further investigations of potent, and useless, pathogen strains will unveil the real potential of boosted-SIT.

## Funding

We are grateful to Sunday Ekesi and Ivan Rwomushana for enabling this project with ICIPE. This work received financial support from a Long-term EU- Africa research and innovation Partnership on food and nutrition security and sustainable Agriculture (LEAP-Agri), project Pest-Free Fruit, in the framework of the European Union’s Horizon 2020 research and innovation programme under grant agreement No 727715, the French Agence Nationale de la Recherche (ANR-18-LEAP-0006-02), and the ECOPHYTO scheme through the grant ANR SUZUKIISS:ME (ANR-21-ECOM-0002).

## Conflicts of interest

The authors declare no conflicts of interest.

## Notes

### Competing Interest Statement

The authors have declared no competing interest.

