## Supplementary material 1 for "Sex-specific sub-lethal effects of low virulence entomopathogenic fungi may boost the Sterile Insect Technique": Duneau_BioRxiv_SuppMat_1.html


Code 

- Show All Code
- Hide All Code

### Sex-specific sub-lethal effects may boost the Sterile Insect Technique with low virulence entomopathogenic agents.

###### by David Duneau

###### February 2024

### Library, Data import and reformatting

```
library(scales)
library(kableExtra)
library(multcomp)
library(lmtest)
library(multcomp)
library(spaMM)
library("survminer")
library(dplyr)
library(tidyr)
library(spaMM)
library(gridExtra)
library(grid)
library(gtable)
library(sjPlot)
library(lme4)
library(car)
library(vcd)
library(visreg)
library(cowplot)
library(ggplotify)
library(gridGraphics)
library(data.table)
library(stringr)
library(rsq)
library(tibble)
library(boot.pval)
library(forcats)
library(ggrepel)
library(coxme)
library(multcomp)
library(dunn.test)

grab_grob <- function(){
  grid.echo()
  grid.grab()
}

RIGHT = function(x,n){
  substring(x,nchar(x)-n+1)
}

LEFT = function(x,n){
  substring(x,1,nchar(x)-n+1)
}

r2.corr.mer <- function(m) {
  lmfit <-  lm(model.response(model.frame(m)) ~ fitted(m))
  summary(lmfit)$r.squared
}

logit2prob <- function(logit){
  odds <- exp(logit)
  prob <- odds / (1 + odds)
  return(prob)
}

`%notin%` = Negate(`%in%`)

ggplotprep2 <- function(x, times){
  #spreading the surfit dataframe into dataframe per day. 
  d <- data.frame(condition=rep(names(x$strata), x$strata), time=x$time, survival=x$surv, upper=x$upper, lower=x$lower)
  # function to add time point 0 
  fillup0 <- function(s) rbind(c(condition=s, time=0, survival=1, upper=1, lower=1), d[d$condition==s, ], deparse.level = 0)
  
  # function to determine the missing time points
  indexes <- function(x, time) {
    if(x%in%time) return(x)
    return(time[which.min(abs(time[time<x]-x))])
  }
  #Function to complete the missing time points
  fillup <- function(s) {
    d.temp <- d[d$condition==s, ]
    time <- as.numeric(d.temp$time)
    id <- sapply(times, indexes, time=time)
    d.temp <- d.temp[match(id, time), ]
    d.temp$time <- times
    return(d.temp)
  }
  
  if(times[1]==0) d <- do.call("rbind", sapply(names(x$strata), fillup0, simplify=F))
  d <- do.call("rbind", sapply(names(x$strata), fillup, simplify=F))
  d <- data.frame(Condition=d$condition, Time=as.numeric(d$time), Survival=as.numeric(d$survival), Upper=as.numeric(d$upper), Lower=as.numeric(d$lower))
  return(d)
} 

extract_coxme_table = function (mod){
    beta = fixef(mod)
    nvar = length(beta)
    nfrail = nrow(mod$var) - nvar
    se = sqrt(diag(mod$var)[nfrail + 1:nvar])
    z= round(beta/se, 2)
    p= format(as.numeric(pchisq((beta/se)^2, 1,lower.tail = F)), 4)
    table=data.frame(cbind(beta,se,z,p))
    return(table)
 }
```

```
SuperSmallfont= 6
Smallfont= 12
Mediumfont= 14
Largefont= 14
verylargefont = 16
pointsize= 0.7
linesize=0.35
meansize = 1.5
Margin=c(0,0,0,0)

fontsizeaxes = 12
fontsizeaxes2 = 10


basic_theme_plot=
  theme(axis.title.x = element_text(size=Mediumfont),
        axis.title.y = element_text(size=Mediumfont),
        plot.caption =element_text(size=Mediumfont,face="italic"),
        axis.line.x = element_line(colour="black",size=0.75),
        axis.line.y = element_line(colour="black",size=0.75),
        axis.ticks.x = element_line(size = 0.75),
        axis.ticks.y = element_line(size = 0.75),
        axis.text.x = element_text(size=Smallfont,colour="black"),
        axis.text.y = element_text(size=Smallfont,colour="black"),
        plot.margin = unit(Margin, "cm"),
        strip.text.x = element_text(size =Mediumfont, colour = "black",face="italic"),
        strip.text.y = element_text(size =Mediumfont, colour = "black",face="italic"),
        strip.background = element_rect(fill=NA, colour="black"),
        strip.placement="outside",
        panel.background = element_blank(),
        legend.key.height = unit(0.4, "cm"),
        legend.key.width= unit(0.6, "cm"),
        legend.title = element_text(face="italic",size=Smallfont), 
        legend.key = element_rect(colour = 'white', fill = "white", linetype='dashed'),
        legend.text = element_text(size=Smallfont),
        legend.background = element_rect(fill=NA))

basic_theme_Surv=
  theme(aspect.ratio = 1,
        panel.background = element_blank(),
        plot.caption =element_text(size=Mediumfont,face="italic"),
        title =element_text(size=Smallfont),
        strip.text.x = element_text(size =Mediumfont, colour = "black",face="italic"),
        strip.text.y = element_text(size =Mediumfont, colour = "black",face="italic"),
        strip.background = element_rect(fill=NA, colour="black"),
        strip.placement="outside",
        axis.title.x = element_text(size=Mediumfont,colour="black"),
        axis.title.y = element_text(size=Mediumfont,colour="black"), 
        axis.line.x = element_line(colour="black",size=0.75),
        axis.line.y = element_line(colour="black",size=0.75),
        axis.ticks.x = element_line(size = 0.75),
        axis.ticks.y = element_line(size = 0.75),
        axis.text.x = element_text(size=Smallfont,colour="black"),
        axis.text.y = element_text(size=Smallfont,colour="black"),
        plot.margin = unit(Margin, "cm"),
        legend.direction = "horizontal", 
        legend.box = "horizontal",
        legend.position = "top",
        legend.key.height = unit(0.4, "cm"),
        legend.key.width= unit(0.6, "cm"),
        legend.title = element_text(face="italic",size=Smallfont), 
        legend.key = element_rect(colour = 'white', fill = "white", linetype='dashed'),
        legend.text = element_text(size=Smallfont),
        legend.background = element_rect(fill=NA))
```

```
#initialize three empty lists for storing the data, file paths, and models, respectively.
d <- list()
path <- list()

# through each CSV file in the specified directory and its subdirectories.
for(f in list.files(path=path.to.data,pattern="*.csv$",recursive=T,full.names=T)) {
  nom <- gsub(".*/(.*).csv","\\1",f)    #extract the file name without the extension.
  cat(nom,"\n") #show the name
  path[[nom]] <- gsub("(.*)/.*csv","\\1/",f) #extract the path to the directory containing the file.
  d[[nom]] <- read.table(f,header=T,sep=";",dec=",")# read in the CSV file and store it in the d list under the corresponding file name.
  
}

#create new data frames by selecting the data stored in the d list under the corresponding file names.
df_inoculum = d[["Spore_counts_2020_06_29"]] %>% 
  mutate(Replicate=paste0("Rep",Replicate),
         Strain=str_remove_all(Strain," "),
         Sample_ID=paste(Strain,Replicate,sep="_"),
         across(where(is.character),as.factor)) %>% 
  relocate(Spore.concentration,.after=Sample_ID) %>% 
  select(-Column_1)

df_surv= d[["Survival_from_overview_2020_01_28_corrected_sf"]] %>% 
  mutate(replicate_ID=str_remove_all(replicate_ID,pattern = "licate ")) %>% 
  select(-TRT_ID_with_replicate) %>%
  mutate(ID = row_number()) %>%
  pivot_longer(cols = starts_with("male_mortality"), names_to = "Time_post_inoculum", values_to = "Deaths") %>%
  uncount(weights = Deaths) %>%
  mutate(Time_post_inoculum=str_remove_all(Time_post_inoculum, "male_mortality_day_"),
         Censor=ifelse(Time_post_inoculum=="male_mortality_Number_that_did_not_die",0,1),
         Time_post_inoculum=str_replace_all(Time_post_inoculum, "male_mortality_Number_that_did_not_die","7"),
         Bloc=str_remove_all(Bloc,"\\?"),
         Time_post_inoculum = as.numeric(Time_post_inoculum),
         across(where(is.character),as.factor)) %>% 
  data.frame()

df_egg = d[["Fecundity_from_overview_stacked_2020_01_28_SF"]] %>% 
  select(-c(Log_egg_numbers)) %>% 
  mutate(bloc=str_remove_all(bloc,"\\?"),
         Cage_ID = str_replace_all(Cage_ID, "-", "_"),
         Fungal_treatment = str_remove_all(Fungal_treatment, " "),
         Treatment=ifelse(Fungal_treatment=="control","Control","Fungus"),
         across(where(is.character),as.factor))
```

### 1 **Spore inoculum (Figure 1)**

Short methods:

We estimated the spore inoculum for each strains.

---

#### What is the spore inoculum for each strain?

---

```
df_inoculum= df_inoculum %>% mutate(Log2_spore=log2(Spore.concentration))
  
model_full_gaussian = fitme( Log2_spore ~ Strain, family= gaussian(), data = df_inoculum)

ci = list()
for (i in c(1:8)){
  ci[[i]] = confint(model_full_gaussian,parm=function(v) fixef(v)[i], boot_args=list(nsim=99, seed=123))
}

df_Confint_inoc = data.frame(est=sapply(ci,function(z) z$t0),
                             inf=sapply(ci,function(z) z$normal[2]),
                             sup=sapply(ci,function(z) z$normal[3]))%>% 
  mutate(est = ifelse(row_number() %in% c(2:8), est + est[1], est),
         inf = ifelse(row_number() %in% c(2:8), inf + est[1], inf),
         sup = ifelse(row_number() %in% c(2:8), sup + est[1], sup)) %>% 
  rownames_to_column("Strain") %>% 
  mutate(Strain=str_remove_all(Strain,"Strain"),
         Strain=str_replace_all(Strain,"Intercept","ICIPE07"),
         Strain=str_remove_all(Strain,"\\("),
         Strain=str_remove_all(Strain,"\\)"))

kruskal = kruskal.test(log2(Spore.concentration) ~ Strain, data = df_inoculum)
```

Males were inoculated with approximately 250-1000 spores (i.e.,
between 2^8 and 2^10 spores). The 95% confidence intervals for all
strains overlapped with each other, suggesting that there is no
significant differences among the strains for the inoculum
(Kruskal-Wallis test: df = 7, Chi-squared = 11.4, p = 0.12). Red dots
and error bars represent the means of spore inoculum and their 95%
confidence intervals estimated by bootstraps.  
(ndlr: in the word document, the inoculum seem to be higher.)

### 2 **Survival (Figure 2)**

Short methods:

We monitored the survival of male and female flies for a duration of
7 days. Males were exposed to spores from various strains (n=7) through
inoculation, while females were exposed through contact with these
males. Each replicate comprised 10 males and 20 females housed in the
same cage. We conducted three replicates on the same day, involving 2 or
3 strains. It is important to note that the strains were not replicated
across multiple days. For each experimental day, we documented the
survival of males and females that were not exposed to the fungus.

---

#### Is there a baseline difference in mortality over a span of 7 days between males and females, as well as among different experiments?

---

There were no significant differences in baseline mortality, observed
when individuals were not exposed to spores, between sexes and across
different exposure days.

| coxme(Surv(Time\_post\_inoculum, Censor)~ Sex \* Bloc + (1\|Cage\_ID) | | | |
| --- | --- | --- | --- |
|  | df | Chi2 | p-value |
| Sex | 1 | 3.17 | 0.075 |
| Day of inoculum | 2 | 2.47 | 0.291 |
| Sex x Day | 2 | 0.07 | 0.965 |

#### How susceptible are males and females to each strain?

```
survdata = survfit(Surv(Time_post_inoculum, Censor)~  Fungal_treatment + Sex , data=df_surv)

toplot = ggplotprep2(survdata, times=c(seq(0,7,by=1))) %>%
  mutate(Condition=str_remove_all(Condition,"[a-zA-Z]*="))%>% 
  separate(Condition, c("Fungal_treatment","Sex"),sep=", ") %>% 
  mutate(Fungal_treatment=str_remove(Fungal_treatment, "Fungal_"),
         Fungal_treatment=str_remove_all(Fungal_treatment, " "),
         Sex=str_remove_all(Sex, " "),
         Control=ifelse(Fungal_treatment=="control","Control","Fungus"),
         across(where(is.character),as.factor))

(Plot_susceptibility = 
    ggplot(toplot, aes(x=Time,y=Survival, group=Fungal_treatment))+
   labs(caption = "(Figure 2A)")+
    facet_grid(.~Sex)+
    geom_vline(xintercept = 2, linetype="dotted")+
    geom_line(aes(linetype=Control,colour=Fungal_treatment),linewidth=1)+
    geom_point(colour="black",size = 1) +
 #   geom_errorbar(data=subset(toplot,Time==4),aes(ymin=Lower, ymax=Upper), width=.1, alpha=0.4, size=1, show.legend=FALSE)+
    scale_color_manual(values=c("black","red","blue","green","darkblue","orange","brown","cyan","darkorange"))+
    scale_linetype_manual(values=c("solid","dashed"))+
    scale_y_continuous("Proportion of survivors",
                       limits=c(0, 1),breaks=c(0,0.2,0.4,0.6,0.8,1))+
    scale_x_continuous("Time since beginning of the experiment",
                       breaks=c(seq(0,7,by=1)))+
    basic_theme_Surv+
    theme(legend.position = "right",
          legend.box = "vertical",
          legend.direction = "vertical"))
```

Both male and female individuals exhibit clear mortality due to the
infection. Females demonstrate mortality approximately 1 day after
males, suggesting a lag associated with transmission from males to
females. Males, who were directly inoculated with spores, experienced
higher mortality rates from the infection compared to females exposed to
infected males through contact. All male individuals succumbed to the
infection 6 days after inoculation, whereas more than 50% of the females
remain alive even after 7 days from the start of the experiments. We
cannot exclude at this point that females which survived were exposed to
the spores.

---

#### Is there a difference among fungal strains?

```
tab_HR=NULL
tab_res_surv_F=NULL
tab_res_surv_M=NULL

tmp=subset(df_surv, Fungal_treatment!="control") %>% 
  mutate(Fungal_treatment = fct_relevel(Fungal_treatment,"ICIPE69"),
         Sex2=fct_relevel(Sex,"Males"),
         Cage_ID=paste(Fungal_treatment, replicate_ID, Sex, sep="_")) %>% 
  droplevels()

model_Surv_M= with(tmp,
                   coxme(Surv(Time_post_inoculum, Censor)~ Fungal_treatment + Fungal_treatment : Sex2 + (1|Cage_ID)))

model_Surv_F= with(tmp,
                   coxme(Surv(Time_post_inoculum, Censor)~ Fungal_treatment + Fungal_treatment : Sex + (1|Cage_ID)))

Confint_M = as.data.frame(confint(model_Surv_M, R=999, level=0.95)) %>% 
  rownames_to_column(var="Strain")
Confint_F = as.data.frame(confint(model_Surv_F, R=999, level=0.95)) %>% 
  rownames_to_column(var="Strain")

tab_res_surv_F = extract_coxme_table(model_Surv_F) 
tab_res_surv_F = tab_res_surv_F %>% 
  mutate(Strain=rownames(tab_res_surv_F))

tab_res_surv_F = left_join(tab_res_surv_F,Confint_F)
  tab_res_surv_F = tab_res_surv_F %>% 
  mutate(Strain=str_remove(Strain,"Fungal_treatment"),
         Sex="Female")%>% 
  slice(1:7)

tab_res_surv_M = extract_coxme_table(model_Surv_M) 
tab_res_surv_M =
  tab_res_surv_M %>% 
  mutate(Strain=rownames(tab_res_surv_M))
tab_res_surv_M = left_join(tab_res_surv_M,Confint_M)
  tab_res_surv_M = tab_res_surv_M %>% 
  mutate(Strain=str_remove(Strain,"Fungal_treatment"),
         Sex="Male")%>% 
  slice(1:7)

tab_HR = rbind(tab_res_surv_M,tab_res_surv_F)%>%
  relocate(Strain, .before=beta)%>%
  relocate(Sex, .before=beta)
colnames(tab_HR) = c("Strain","Sex","beta","se","z","p", "beta_lower", "beta_upper")

# tab_HR_early_infection = left_join(tab_HR_early_infection,d_Block_surv)
# tab_HR_early_infection$Block= as.character(tab_HR_early_infection$Block)
tab_HR[nrow(tab_HR)+1,] = c("ICIPE69","Male",0, 10^-6, rep(NA,2),0,0)
tab_HR[nrow(tab_HR)+1,] = c("ICIPE69","Female",0, 10^-6, rep(NA,2),0,0)

tab_HR = 
  tab_HR%>%
  mutate(beta=as.numeric(as.character(beta)),
         HazardRatio = exp(beta),
         se = as.numeric(as.character(se)),
         z = as.numeric(as.character(z)),
         p = as.numeric(as.character(p)),
         beta_upper = as.numeric(as.character(beta_upper)),
         beta_lower = as.numeric(as.character(beta_lower)),
         across(where(is.character),factor))

##### Susceptibility in female ####
tab_F = subset(tab_HR,Sex=="Female")%>%
  droplevels() %>% 
  mutate(Strain=reorder(Strain,desc(beta)),
         across(where(is.character),as.factor))

tmp=select(tab_F,c(1:2,7,8))
Sample_size_F=
  subset(df_surv, Sex=="Females")%>%
  group_by(Fungal_treatment,Sex)%>%
  summarise(Sample_size=n())%>%
  mutate(Fungal_treatment = factor(Fungal_treatment, levels=c(levels(tab_F$Strain))))

Sample_size_F = left_join(Sample_size_F,tmp, by=c("Fungal_treatment"="Strain"))  %>% 
  mutate(Fungal_treatment=fct_relevel(Fungal_treatment,levels(tab_F$Strain)))

Sample_size_F_1 =  Sample_size_F %>% 
  filter(Fungal_treatment %in% c(levels(tab_F$Strain)[seq(0,nrow(Sample_size_F),by=2)]))  %>% 
  dplyr::rename("Strain"="Fungal_treatment")%>%
  mutate(Strain=fct_relevel(Strain,levels(tab_F$Strain)))%>% 
  data.frame()

Sample_size_F_2 =  Sample_size_F %>% 
  filter(Fungal_treatment %in% c(levels(tab_F$Strain)[seq(1,nrow(Sample_size_F),by=2)]))  %>% 
  filter(!is.na(Fungal_treatment)) %>% 
  dplyr::rename("Strain"="Fungal_treatment")%>%
  mutate(Strain=fct_relevel(Strain,levels(tab_F$Strain)))%>% 
  data.frame()

plot_HR_F=
    ggplot(tab_F, aes(x=Strain, y=beta)) + 
    facet_grid(.~Sex)+
    geom_errorbar(aes(ymin=beta_lower, ymax=beta_upper), col="black",width=.17,  size=0.5, show.legend=FALSE)+
    geom_point( stat="identity", show.legend=FALSE,  size=1.5)+
    geom_hline(yintercept = 0,colour="red",linetype=4)+
    scale_color_manual(values=c("red","blue","green","orange","black"))+
    scale_y_continuous("log(HR) (95 % CI)\nrelative to ICIPE69",
                       breaks=c(seq(-5,5,by=1)))+
    coord_cartesian(ylim = c(-2, 2), expand = T, clip = "on")+
    geom_text(data = Sample_size_F_1, mapping = aes(x = Strain, y = beta_lower-0.4, label = paste("(",Sample_size,")",sep="")),color="orange",size=2.7)+
    geom_text(data = Sample_size_F_2, mapping = aes(x = Strain, y = beta_upper+0.4 , label = paste("(",Sample_size,")",sep="")),color="orange",size=2.7)+
    scale_x_discrete("Strain")+
    basic_theme_plot+
    theme(axis.text.x =element_text(angle=45,hjust=1),
          aspect.ratio = 0.5)

##### Susceptibility in male ####
tab_M = subset(tab_HR,Sex=="Male")%>%
  droplevels() %>% 
  mutate(Strain=reorder(Strain,desc(beta)),
         across(where(is.character),as.factor)) %>% 
  data.frame()

tmp=select(tab_M,c(1:2,7,8))
Sample_size_M=
  subset(df_surv, Sex=="Males")%>%
  group_by(Fungal_treatment,Sex)%>%
  summarise(Sample_size=n())%>%
  mutate(Fungal_treatment = factor(Fungal_treatment, levels=c(levels(tab_M$Strain))))

Sample_size_M = left_join(Sample_size_M,tmp, by=c("Fungal_treatment"="Strain"))  %>% 
  mutate(Fungal_treatment=fct_relevel(Fungal_treatment,levels(tab_M$Strain)))

Sample_size_M_1 =  Sample_size_M %>% 
  filter(Fungal_treatment %in% c(levels(tab_M$Strain)[seq(0,nrow(Sample_size_M),by=2)]))  %>% 
  dplyr::rename("Strain"="Fungal_treatment")%>%
  mutate(Strain=fct_relevel(Strain,levels(tab_M$Strain)))%>% 
  data.frame()

Sample_size_M_2 =  Sample_size_M %>% 
  filter(Fungal_treatment %in% c(levels(tab_M$Strain)[seq(1,nrow(Sample_size_M),by=2)]))  %>% 
  filter(!is.na(Fungal_treatment)) %>% 
  dplyr::rename("Strain"="Fungal_treatment")%>%
  mutate(Strain=fct_relevel(Strain,levels(tab_M$Strain)))%>% 
  data.frame()

plot_HR_M=
    ggplot(tab_M, aes(x=Strain, y=beta)) + 
    facet_grid(.~Sex)+
    geom_errorbar(aes(ymin=beta_lower, ymax=beta_upper), col="black",width=.17,  size=0.5, show.legend=FALSE)+
    geom_point( stat="identity", show.legend=FALSE,  size=1.5)+
    geom_hline(yintercept = 0,colour="red",linetype=4)+
    scale_color_manual(values=c("red","blue","green","orange","black"))+
    scale_y_continuous("",
                       breaks=c(seq(-5,5,by=1)))+
    coord_cartesian(ylim = c(-2, 2), expand = T, clip = "on")+
    geom_text(data = Sample_size_M_1, mapping = aes(x = Strain, y = beta_lower-0.4, label = paste("(",Sample_size,")",sep="")),color="orange",size=2.7)+
    geom_text(data = Sample_size_M_2, mapping = aes(x = Strain, y = beta_upper+0.5 , label = paste("(",Sample_size,")",sep="")),color="orange",size=2.7)+
    scale_x_discrete("Strain")+
    basic_theme_plot+
    theme(axis.text.x =element_text(angle=45,hjust=1),
          aspect.ratio = 0.5)
```

We calculate the log(Hazard ratio) for each strains relative to the
most pathogenic strains in males (i.e. ICIPE69). The log hazard ratio
represents the logarithm of the ratio of hazards between the reference
strain and the other strains. It quantifies the relative difference in
the risk of dying among strains.

No significant difference in female mortality was observed between
strains. However, strains ICIPE07 and ICIPE69 exhibited a higher rate of
male mortality compared to other strains, indicating their lower
suitability, particularly when considering strain ICIPE41, for Sterile
Insect Technique. It is important to note, however, that the most
virulent strains were tested on the same day (Supplementary figure 1).
While the baseline mortality on that day was not higher, it is worth
considering that the higher mortality could potentially be influenced by
environmental conditions specific to that day.

```
survdata = survfit(Surv(Time_post_inoculum, Censor)~ Bloc + Fungal_treatment + Sex , data=df_surv)

toplot = ggplotprep2(survdata, times=c(seq(0,7,by=1))) %>%
  mutate(Condition=str_remove_all(Condition,"[a-zA-Z]*="))%>% 
  separate(Condition, c("Block","Fungal_treatment","Sex"),sep=", ") %>% 
  mutate(Fungal_treatment=str_remove(Fungal_treatment, "Fungal_"),
         Fungal_treatment=str_remove_all(Fungal_treatment, " "),
         Sex=str_remove_all(Sex, " "),
         Control=ifelse(Fungal_treatment=="control","Control","Fungus"),
         Block= factor(Block,c("13/05/2019","28/05/2019","11/06/2019")),
         across(where(is.character),as.factor))

(Plot_susceptibility_block = 
    ggplot(toplot, aes(x=Time,y=Survival, group=Fungal_treatment))+
    geom_vline(xintercept = 2, linetype="dotted")+
   labs(caption = "(Figure Supp1)")+
    facet_grid(Block~Sex)+
    geom_line(aes(linetype=Control,colour=Fungal_treatment),linewidth=1)+
    geom_point(aes(colour=Fungal_treatment),size = pointsize) +
    geom_errorbar(data=subset(toplot,Time==5),aes(ymin=Lower, ymax=Upper), width=.1, alpha=0.4, size=1, show.legend=FALSE)+
    scale_color_manual("Treatment",values=c("black","red","blue","green","darkblue","orange","brown","cyan","darkorange"))+
    scale_linetype_manual(values=c("solid","dotted"))+
    scale_y_continuous("Proportion of survivors",
                       limits=c(0, 1),breaks=c(0,0.2,0.4,0.6,0.8,1))+
    scale_x_continuous("Time since beginning of the experiment",
                       breaks=c(seq(0,7,by=1)))+
    basic_theme_plot+
    theme(legend.position = "right",
          legend.box = "vertical",
          legend.direction = "vertical"))
```

#### Is there a correlation between inoculum and male mortality?

```
tab_HR_sub = tab_HR %>% 
  filter(Sex=="Male") %>% 
  select(c(Strain,beta,beta_lower,beta_upper))

tab_cor_inoc_HR = left_join(tab_HR_sub,df_Confint_inoc)

correlation.test = cor.test(tab_cor_inoc_HR$beta,tab_cor_inoc_HR$est,method = "pearson")
```

As anticipated due to the lack of statistical differences in spore
inoculum among strains, there was no correlation observed between the
inoculum of spores in males and their survival. Furthermore, the
tendency leaned towards a negative correlation between inoculum and
survival (Pearson correlation test: df = 6, correlation = -0.33, p =
0.4). Taken together, our result indicates that the variation in
virulence is likely attributed to the pathogenicity of the strain.

### 3 **Egg laying (Figure 3)**

Short methods:

During the 7-day duration of the experiment, we observed the
fecundity of female flies on mango skin. At the beginning of the
experiment, each cage contained 20 females, and their numbers gradually
decreased throughout the study period as they die. As a result, the egg
count was not based on a constant number of females.

---

#### Is there a decrease in egg laying due to exposure to the fungus?

---

```
df_egg_mean =
  df_egg %>% 
  group_by(bloc,Fungal_treatment,Treatment,Cage_ID) %>% 
  summarise(Total_egg=sum(Egg_numbers)) %>% 
  data.frame() %>% 
  mutate(bloc=factor(bloc,c("13/05/2019","28/05/2019","11/06/2019")))


tmp= subset(df_egg_mean, bloc=="11/06/2019")
kruskal_1 = kruskal.test(log2(Total_egg) ~ Fungal_treatment , data = tmp)

dunn_test_1 = dunn.test(tmp$Total_egg,tmp$Fungal_treatment, method = "bh") %>% 
  as.data.frame() %>% 
  relocate(comparisons, .before=chi2) %>% 
  arrange(P.adjusted)

tmp= subset(df_egg_mean, bloc=="13/05/2019")
kruskal_2 = kruskal.test(log2(Total_egg) ~ Fungal_treatment , data = tmp)

dunn_test_2 = dunn.test(tmp$Total_egg,tmp$Fungal_treatment, method = "bh") %>% 
  as.data.frame() %>% 
  relocate(comparisons, .before=chi2) %>% 
  arrange(P.adjusted)

tmp= subset(df_egg_mean, bloc=="28/05/2019")
kruskal_3 = kruskal.test(log2(Total_egg) ~ Fungal_treatment , data = tmp)

dunn_test_3 = dunn.test(tmp$Total_egg,tmp$Fungal_treatment, method = "bh") %>% 
  as.data.frame() %>% 
  relocate(comparisons, .before=chi2)%>% 
  arrange(P.adjusted)
```

We observed a significant difference in fecundity among treatments
(Figure 3A). The total number of eggs per cage was monitored over a span
of 7 days in three separate experiments. Given our experimental design
with a limited sample size of three cages per treatment and the need to
employ non-parametric statistical tests, the letters in the figure
represent the statistical differences after p-value corrections with the
probability to falsely attribute a difference (error type I) of 10%,
instead of 5%.

There was more eggs in the control treatment of the third day of
experiments, likely because we provided two halves of mango skin in all
treatments done this day. Competition for space on the mango skin may
have affected quantitatively the fecundity on the other days. However,
on day 13/05, the control had a lower fecundity than the fungal
treatments, suggesting that the control treatment were not competiting
for space. On day 28/05, the control had a significantly higher
fecundity, therefore, the effect would only be stronger.

In general, most strains resulted in a decrease in the number of eggs
per cage, indicating a potential impact on fecundity. However, this
reduction of fecundity could be due to female mortality within the
cages. Interestingly, strains ICIPE69, and ICIPE07 exhibited contrasting
behavior, as they showed higher fecundity compared to the control
despite their high virulence. We nevertheless decided to test the
difference of egg number per cage within the 2 days after females were
in contact with males.

#### Is there a decrease in egg laying due to exposure to the fungus within the 2 first days after inoculation?

```
df_egg_mean_2d =
  df_egg %>% 
  filter(Time_since_spore_exposure<=2) %>% 
  group_by(bloc,Fungal_treatment,Treatment,Cage_ID) %>% 
  summarise(Total_egg=sum(Egg_numbers)) %>% 
  data.frame() %>% 
  mutate(bloc=factor(bloc,c("13/05/2019","28/05/2019","11/06/2019")))

tmp= subset(df_egg_mean_2d, bloc=="11/06/2019")
kruskal_4 = kruskal.test(log2(Total_egg) ~ Fungal_treatment , data = tmp)

dunn_test_4 = dunn.test(tmp$Total_egg,tmp$Fungal_treatment, method = "bh") %>% 
  as.data.frame() %>% 
  relocate(comparisons, .before=chi2) %>% 
  arrange(P.adjusted)

tmp= subset(df_egg_mean_2d, bloc=="13/05/2019")
kruskal_5 = kruskal.test(log2(Total_egg) ~ Fungal_treatment , data = tmp)

dunn_test_5 = dunn.test(tmp$Total_egg,tmp$Fungal_treatment, method = "bh") %>% 
  as.data.frame() %>% 
  relocate(comparisons, .before=chi2) %>% 
  arrange(P.adjusted)

tmp= subset(df_egg_mean_2d, bloc=="28/05/2019")
kruskal_6 = kruskal.test(log2(Total_egg) ~ Fungal_treatment , data = tmp)

dunn_test_6 = dunn.test(tmp$Total_egg,tmp$Fungal_treatment, method = "bh") %>% 
  as.data.frame() %>% 
  relocate(comparisons, .before=chi2)%>% 
  arrange(P.adjusted)
```

We analysed the total number of eggs per cage over a span of 2 days
in three separate experiments. Two days corresponds to the period were
females did not die yet. The pattern did not differ strongly with the
total number of eggs per cage over a span of 7 days, suggesting that the
difference in number of eggs was not driven by the reduction in number
of females per cage over time due to mortality.

---

Tables accompanying figure 3A.

```
dunn_test_1 = data.frame(lapply(dunn_test_1, function(x) {
  if(is.numeric(x)) signif(x, digits = 2)
  else x
}))

dunn_test_2 = data.frame(lapply(dunn_test_2, function(x) {
  if(is.numeric(x)) signif(x, digits = 2)
  else x
}))

dunn_test_3 = data.frame(lapply(dunn_test_3, function(x) {
  if(is.numeric(x)) signif(x, digits = 2)
  else x
}))


dunn_test_1%>%
  kable(col.names = c("Comparisons Dunn test","Chi2" ,"Z", "p-value", "p-value adj (FDR)"),row.names = FALSE) %>%   
  add_header_above(c("Fecundity over 7 days; 11/06/2019; kruskal-wallis p = 0.04" = 5))%>%
  kable_styling(bootstrap_options = c("striped", "hover", "condensed"), full_width = F)
```

| Fecundity over 7 days; 11/06/2019; kruskal-wallis p = 0.04 | | | | |
| --- | --- | --- | --- | --- |
| Comparisons Dunn test | Chi2 | Z | p-value | p-value adj (FDR) |
| control - ICIPE387 | 8.3 | 1.90 | 0.027 | 0.041 |
| control - ICIPE20 | 8.3 | 2.00 | 0.021 | 0.042 |
| ICIPE387 - ICIPE62 | 8.3 | -2.00 | 0.021 | 0.062 |
| ICIPE20 - ICIPE62 | 8.3 | -2.20 | 0.016 | 0.094 |
| control - ICIPE62 | 8.3 | -0.11 | 0.450 | 0.450 |
| ICIPE20 - ICIPE387 | 8.3 | -0.11 | 0.450 | 0.550 |

```
dunn_test_2%>%
  kable(col.names = c("Comparisons Dunn test","Chi2" ,"Z", "p-value", "p-value adj (FDR)"),row.names = FALSE) %>%   
  add_header_above(c("Fecundity over 7 days; 13/05/2019; kruskal-wallis p = 0.06" = 5))%>%
  kable_styling(bootstrap_options = c("striped", "hover", "condensed"), full_width = F)
```

| Fecundity over 7 days; 13/05/2019; kruskal-wallis p = 0.06 | | | | |
| --- | --- | --- | --- | --- |
| Comparisons Dunn test | Chi2 | Z | p-value | p-value adj (FDR) |
| control - ICIPE69 | 5.6 | -2.20 | 0.013 | 0.038 |
| control - ICIPE07 | 5.6 | -1.80 | 0.037 | 0.055 |
| ICIPE07 - ICIPE69 | 5.6 | -0.45 | 0.330 | 0.330 |

```
dunn_test_3%>%
  kable(col.names = c("Comparisons Dunn test","Chi2" ,"Z", "p-value", "p-value adj (FDR)"),row.names = FALSE) %>%   
  add_header_above(c("Fecundity over 7 days; 28/05/2019; kruskal-wallis p = 0.09" = 5))%>%
  kable_styling(bootstrap_options = c("striped", "hover", "condensed"), full_width = F)
```

| Fecundity over 7 days; 28/05/2019; kruskal-wallis p = 0.09 | | | | |
| --- | --- | --- | --- | --- |
| Comparisons Dunn test | Chi2 | Z | p-value | p-value adj (FDR) |
| control - ICIPE18 | 6.4 | 1.90 | 0.027 | 0.054 |
| control - ICIPE402 | 6.4 | 2.30 | 0.012 | 0.071 |
| control - ICIPE41 | 6.4 | 1.90 | 0.027 | 0.081 |
| ICIPE18 - ICIPE402 | 6.4 | 0.34 | 0.370 | 0.440 |
| ICIPE18 - ICIPE41 | 6.4 | 0.00 | 0.500 | 0.500 |
| ICIPE402 - ICIPE41 | 6.4 | -0.34 | 0.370 | 0.550 |

---

Tables accompanying figure 3B.

```
dunn_test_4 = data.frame(lapply(dunn_test_4, function(x) {
  if(is.numeric(x)) signif(x, digits = 2)
  else x
}))

dunn_test_5 = data.frame(lapply(dunn_test_5, function(x) {
  if(is.numeric(x)) signif(x, digits = 2)
  else x
}))

dunn_test_6 = data.frame(lapply(dunn_test_6, function(x) {
  if(is.numeric(x)) signif(x, digits = 2)
  else x
}))


dunn_test_4%>%
  kable(col.names = c("Comparisons Dunn test","Chi2" ,"Z", "p-value", "p-value adj (FDR)"),row.names = FALSE) %>%   
  add_header_above(c("Fecundity over 2 days; 11/06/2019; kruskal-wallis p = 0.08" = 5))%>%
  kable_styling(bootstrap_options = c("striped", "hover", "condensed"), full_width = F)
```

| Fecundity over 2 days; 11/06/2019; kruskal-wallis p = 0.08 | | | | |
| --- | --- | --- | --- | --- |
| Comparisons Dunn test | Chi2 | Z | p-value | p-value adj (FDR) |
| ICIPE387 - ICIPE62 | 6.9 | -2.00 | 0.021 | 0.062 |
| ICIPE20 - ICIPE62 | 6.9 | -2.20 | 0.016 | 0.094 |
| control - ICIPE387 | 6.9 | 1.50 | 0.071 | 0.110 |
| control - ICIPE20 | 6.9 | 1.60 | 0.056 | 0.110 |
| control - ICIPE62 | 6.9 | -0.57 | 0.290 | 0.340 |
| ICIPE20 - ICIPE387 | 6.9 | -0.11 | 0.450 | 0.450 |

```
dunn_test_5%>%
  kable(col.names = c("Comparisons Dunn test","Chi2" ,"Z", "p-value", "p-value adj (FDR)"),row.names = FALSE) %>%   
  add_header_above(c("Fecundity over 2 days; 13/05/2019; kruskal-wallis p = 0.19" = 5))%>%
  kable_styling(bootstrap_options = c("striped", "hover", "condensed"), full_width = F)
```

| Fecundity over 2 days; 13/05/2019; kruskal-wallis p = 0.19 | | | | |
| --- | --- | --- | --- | --- |
| Comparisons Dunn test | Chi2 | Z | p-value | p-value adj (FDR) |
| control - ICIPE69 | 3.3 | -1.50 | 0.068 | 0.10 |
| control - ICIPE07 | 3.3 | -1.60 | 0.051 | 0.15 |
| ICIPE07 - ICIPE69 | 3.3 | 0.15 | 0.440 | 0.44 |

```
dunn_test_6%>%
  kable(col.names = c("Comparisons Dunn test","Chi2" ,"Z", "p-value", "p-value adj (FDR)"),row.names = FALSE) %>%   
  add_header_above(c("Fecundity over 2 days; 28/05/2019; kruskal-wallis p = 0.04" = 5))%>%
  kable_styling(bootstrap_options = c("striped", "hover", "condensed"), full_width = F)
```

| Fecundity over 2 days; 28/05/2019; kruskal-wallis p = 0.04 | | | | |
| --- | --- | --- | --- | --- |
| Comparisons Dunn test | Chi2 | Z | p-value | p-value adj (FDR) |
| control - ICIPE41 | 8.1 | 2.70 | 0.0033 | 0.020 |
| control - ICIPE402 | 8.1 | 2.00 | 0.0210 | 0.062 |
| control - ICIPE18 | 8.1 | 1.40 | 0.0870 | 0.130 |
| ICIPE18 - ICIPE41 | 8.1 | 1.40 | 0.0870 | 0.170 |
| ICIPE18 - ICIPE402 | 8.1 | 0.68 | 0.2500 | 0.250 |
| ICIPE402 - ICIPE41 | 8.1 | 0.68 | 0.2500 | 0.300 |

#### Correlation survival and fecundity.

```
tab_HR_sub_F = tab_HR %>% 
  filter(Sex=="Female") %>% 
  select(c(Strain,beta,beta_lower,beta_upper))

mean_fec=
  df_egg_mean %>% 
  group_by(Fungal_treatment) %>% 
  summarize(mean=mean(Total_egg,na.rm=T),
            min=min(Total_egg),
            max=max(Total_egg))


tab_cor_fec_HR = left_join(tab_HR_sub,mean_fec,by=c("Strain"="Fungal_treatment"))

#cor_test_fec_HR = with(tab_cor_fec_HR,
#                       cor.test(beta,mean,method = "pearson"))
```

We analyzed the correlation between survival (log(HR)) and fecundity
(mean) and observed a tendency where strains with lower mortality rates
also exhibited greater reductions in fecundity (Pearson correlation =
0.6, p-value = 0.1).
